# A novel MARV glycoprotein-specific antibody with potentials of broad-spectrum neutralization to filovirus

**DOI:** 10.1101/2023.09.17.558100

**Authors:** Yuting Zhang, Min Zhang, Haiyan Wu, Xinwei Wang, Hang Zheng, Junjuan Feng, Jing Wang, Longlong Luo, He Xiao, Chunxia Qiao, Xinying Li, Yuanqiang Zheng, Weijin Huang, Youchun Wang, Yi Wang, Yanchun Shi, Jiannan Feng, Guojiang Chen

## Abstract

Marburg virus (MARV) is one of the filovirus species that cause deadly hemorrhagic fever in humans, with mortality rates up to 90%. Neutralizing antibodies represent ideal candidates to prevent or treat virus disease. However, no antibody has been approved for MARV treatment to date. In this study, we identified a novel human antibody named AF-03 that targeted MARV glycoprotein (GP). AF-03 possessed a high binding affinity to MARV GP and showed neutralizing and protective activities against the pseudotyped MARV in vitro and in vivo. Epitope identification, including molecular docking and experiment-based analysis of mutated species, revealed that AF-03 recognized the Niemann-Pick C1 (NPC1) binding domain within GP1. Interestingly, we found the neutralizing activity of AF-03 to pseudotyped Ebola viruses (EBOV, SUDV, and BDBV) harboring cleaved GP instead of full-length GP. Furthermore, NPC2-fused AF-03 exhibited neutralizing activity to several filovirus species and EBOV mutants via binding to CI-MPR. In conclusion, this work demonstrates that AF-03 represents a promising therapeutic cargo for filovirus-caused disease.

## Introduction

Filoviruses are a nonsegmented negative-sense RNA viruses, comprised of six genera, Ebolavirus, Marburgvirus, Cuevavirus, Striavirus, Thamnovirus and a recently discovered sixth genus, Dianlovirus^1–3^. The Marburgvirus genus consists of Marburg virus (MARV) and Ravn virus (RAVN)^1,4,5^. The former includes three strains--Uganda, Angola and Musoke. The Ebola virus genus includes six distinct species including Zaire Ebola virus (EBOV), Bundibugyo virus (BDBV), Sudan virus (SUDV), Reston virus (RESTV), Taii Forest virus (TAFV) and Bombali virus (BOMV), the first three of which cause severe hemorrhagic fevers^6,7^. The genus Cuevavirus (Lloviu virus, LLOV) was isolated from Miniopterus schreibersii bats in Spain and Hungary and potently infected monkey and human cells^8,9^. The genus Měnglà virus (MLAV) was discovered in the liver of a bat from Mengla, Yunnan, China in 2019. So far, only an almost complete RNA sequence of the viral genome is available, there are no viable MLAVs isolated.^10^ MARV and EBOV infect humans and non-human primates, causing Marburg virus disease (MVD) and EBOV virus disease (EVD) with an incubation period of 2-21 days^11^. The symptoms of MVD include severe headache and high fever rapidly within 5 days of the onset of symptoms, followed by diarrhea and vomiting, leading to up to 90% fatality rate^11^. Therefore, MARV and EBOV have high potentials to cause a public health emergency.

Glycoprotein (GP) on the surface of filoviruses is a type I transmembrane protein and consists of GP1 and GP2 subunits^12,13^. It is inserted into the virus envelope in the form of homotrimeric spikes^14^ and is responsible for viral attachment and entry. The furin cleaves Marburg GP at the amino acid 435 into two subunits, GP1 and GP2, which remains linked by a disulfide bond^15^. GP1 contains a receptor binding domain (RBD), a glycan cap, and a heavily glycosylated mucin-like domain (MLD), which mediates binding to entry factors and receptors^16^. GP2 has a partial MLD, a transmembrane domain for viral anchoring to the envelop surface, and a fusion peptide required for the fusion of virus and cell membranes^17–19^. In the Ebola virus, the furin cleavage site is located at residue 501 and the entire MLD is attached to the GP1 subunit^20^. Marburg virus contains 66 amino acids on GP2 that are absent from the Ebola virus MLD, and are called “wings” due to their outward projection and flexibility^17^.

Currently, GP is a major target for antibodies validated in filovirus-infected animals and clinical trials because it is exposed on the surface of the virus and plays a key role in viral entry^21^. Filoviruses initially enter cells by endocytosis or macropinocytosis^22,23^. Once inside the endosome, GP is cleaved by host cathepsins and glycan cap and MLD are removed, enabling GP to bind to NPC1 ^24,25^. Interestingly, Ebola viral entry requires cathepsin B cleavage^26^, which is redundant for MARV entry^27,28^. Hashiguchi et al. proposed that the receptor binding domain was masked by glycan cap and MLD in the Ebola virus, whereas it was partially exposed in the Marburg virus^16^.

To date, there is no licensed treatment or vaccine for Marburg infection, although a panel of antibodies with potentials of neutralization has been isolated from a survivor subjected to MARV infection^29^. Herein, we utilized phage display technology to screen an antibody in a well-established antibody library^30,31^ and obtained a novel human antibody with prominent neutralizing activity. Furthermore, NPC2 fusion at the N terminus of the light chain of this antibody potentiates broad-spectrum inhibition of cell entry of filovirus species and mutants.

## Materials and methods

### Cell lines and plasmids

Human embryonic kidney cells HEK293T and human hepatoma cells Huh7 were purchased from ATCC. These cell lines were cultured in Dulbecco’s modified Eagle’s medium (DMEM) (Gibco, 11965e092) supplemented with 100 units/ml penicillin, 100 units/ml streptomycin (Gibco, 15140) and 10% fetal bovine serum (Gibco, 10099) in a humidified atmosphere (5% CO_2_, 95% air) at 37°C. ExpiCHO-S cells were purchased from Gibco and cultured in ExpiCHO™ Expression Medium (Gibco, A29100) in a humidified atmosphere (8% CO_2_, 92% air) on an orbital shaker platform.

MARV (AFV31370.1), Angola (Q1PD50.1), Musoke (YP_001531156.1), RAVN (YP_009055225.1), TAFV (Q66810), RESTV (Q66799), BOMV (YP_009513277.1) MLAV (YP_010087186.1), LLOV (JF828358) GP plasmids were synthesized by GENEWIZ and then cloned into the expression vector pcDNA3.1. EBOV (A0A068J419), BDBV (AYI50382), SUDV (Q7T9D9) GP and HIV-based vector pSG3. Δenv. cmvFluc plasmids were kindly gifted by China Institute for Food and Drug Control.

### Preparation of full-length antibody and antigen

AF-03 was selected from a human phage antibody library, which displays on the surface of M13 bacteriophage particles. Screening procedures were described in detail previously^32,33^. Phage antibodies that bound to MARV GP protein were obtained to express full-length IgG using a standard protocol. In brief, the VH and VL regions of AF-03 were constructed into a mammalian full-length immunoglobulin expression vector pFRT-KIgG1 (Thermo, V601020) to generate plasmid AF-03. The human NPC2 gene (aa20-151) was linked to the VL of AF-03 by a short linker “TVAAP” and then constructed into pFRT-KIgG1 (designated as AF03-NL). The AF-03 and AF03-NL plasmid were transfected into ExpiCHO-S cells using the ExpiFectamine™ CHO Transfection Kit (Gibco, A29129) following the manufacturer’s instructions. Purification was performed using ÄKTA prime plus system (GE Healthcare) with protein A column (GE Healthcare). MARV GP (Uganda strain) (aa 20-648, Δ277-455), CI-MPR1-3 (aa36-466) and NPC2 (aa20-151) gene with six histidine tagged at C-terminus was cloned into mammalian expression vector pcDNA3.1 respectively and then transfected into HEK293T cells. MARV GP was purified using nickel column (GE Healthcare, 11003399). The concentration of proteins and antibodies were quantified by bicinchoninic acid (BCA) method.

### ELISA

The 96-well plates were coated with 2 μg/ml MARV-GP and mutated MARV GP (Q^128^S-N^129^S/C^226^Y) respectively and incubated overnight at 4°C. Wells were washed for three times and blocked for 1 h at 37°C. A series of 12 concentrations of AF-03 and MR78 (starting at 20 μg/ml, 2-fold dilution) were added and incubated for 1 h at 37 °C. Bound antibodies were detected with horseradish peroxidase (HRP)-labeled goat anti-human IgG secondary antibody (Invitrogen, A18817) at 37 °C for 30 min. Binding signals were visualized using a TMB substrate (CWBIO, CW0050S) and the reaction was stopped by adding 2 N H_2_SO_4_. The light absorbance at 450 nm was measured by microplate reader (Thermo Fisher Scientific).

For competitive ELISA, biotinylated AF-03 (1 μg/ml) was coated. MR78 and control mAb (Herceptin) in 3-fold serial dilution (ranging from 200 to 0.27 µg/ml) and added to the plates. After 1 h incubation at 37°C, the plates were washed and the bound biotin-AF-03 was detected by adding horseradish peroxidase (HRP)-labeled Streptavidin (Thermo, S911). After a further 30 min incubation at 37°C, the plates were washed and TMB was added. The reaction was stopped by adding 2 N H_2_SO_4_. Absorbance was measured at 450 nm using a plate reader.

### SPR analysis of antibody affinity

The SPR (surface plasmon resonance) analysis was performed using a Biacore T200 machine with CM5 chips (GE Healthcare) at room temperature (25°C). All the proteins using in SPR analysis were exchanged to BIAcore® buffer, consisting of PBS-P+, with 0.5% Surfactant P20 and 0.5% DMSO, pH 7.4. The chip was subsequently immobilized with MARV GP in sodium acetate, pH 5.0 and then blocked with 1□M ethanolamine, pH 8.0. AF-03 were diluted by running buffer ranging from 0.26 to 0.002 nM. The chip was regenerated with glycine-HCl (pH 2.0, 10 mM). Data were analyzed with Biacore T200 Evaluation Software.

### Computer-guided homology modeling and molecular docking

To uncover the potential epitope of the GP protein, the binding mode between AF-03 and MARV GP protein was analyzed theoretically as following: The three-dimensional theoretical structure of fragment variable (FV) was constructed using computer-guided homology modeling approach (Insight II 2000 sofware, MSI Co., stored in IBM workstation) based on the amino acid sequences of the variable structural domains of the heavy and light chains of AF-03, and the conserved regions (the framework region of the antibody variable domain) and loop structural domains (the CDR region of the antibody variable domain) were identified. The 3-D structure of the AF-03 Fv fragment was optimized under the consistent valence force field (CVFF) using the steepest descent and conjugate gradient minimization methods. The final minimized 3-D structure was evaluated by means of Ramachandran diagrams. In addition, the 3-D theoretical structure of the MARV GP protein was obtained using Alphafold 2 software online and optimized using the CVFF force field in Insight II 2000 software. Under molecular docking method, using the crystal complex structure of MR78 and GP protein (PDB code: 5uqy) as model^16^, the 3-D complex structures AF-03 Fv fragment and GP were obtained and optimized. With the determined 3-D structure of the AF-03 Fv fragment and GP, 50-ns molecular dynamics were performed with the Discovery_3 module. All calculations were performed using Insight II 2000 software (MSI Co., San Diego) with IBM workstation.

### Pseudovirus preparation

HIV vector (pSG3.Δenv.cmvFluc) bearing MARV, mutated MARV (Q^128^S-N^129^S, T^204^A-Q^205^A-T^206^A, Y^218^A, K^222^A and C^226^Y), EBOV (parental and 17 mutants indicated), SUDV, BDBV, TAFV, RESTV, BOMV, RAVN, MLAV and LLOV GPs were prepared by liposome-mediated transfection of HEK293T cells using JetPRIME (Polyplus Transfection, 25Y1801N5) respectively. Cells were seeded in 6-well plates at a density of 7×10^5^cells/well and transfected with 2 μg plasmids (0.4 μg GP and 1.6 μg HIV vector) when cells reached 60-80% confluence. Supernatants were collected 48 h after transfection, centrifuged to remove cell debris at 3,000 rpm for 10 min, filtered through a 0.45 μm-pore filter (Millipore, SLHUR33RB) and stored at -80°C.

### Pseudovirus entry and antibody neutralization assay

Huh7 and HEK293T cells (3×10^4^ cells/100 μl/well) were infected with 100 μl pseudovirus, which contained a luciferase reporter gene respectively. The luciferase activity was measured in a fluorescence microplate reader (Promega). The operation steps were following: after 36 h incubation at 37°C, 100 μl of culture medium were discarded and addition with 100 μl of Bright-Glo luciferase reagent (Promega, E6120) in each well. Mixtures were transferred to 96-well whiteboards after 2-minute reaction to detect the relative luciferase intensity.

For AF-03 neutralization of MARV assays, 50 μl mAb (starting at 20 μg/ml) was 3-fold serially diluted and separately mixed with MARV pseudovirus at the same volume. The mixture was incubated at 37°C for 1 h, followed by the addition of 100 μl cells (3×10^4^ cells/well). 50% of maximal inhibitory concentration was defined as IC_50_. IC_50_ values were determined by non-linear regression with least-squares fit in GraphPad Prism 8 (GraphPad Software).

In terms of AF-03 neutralization of ebola virus assays, pseudovirus (EBOV, SUDV and BDBV) were processed with thermolysin as previously described^34^. Briefly, Pseudotyped ebola virus were incubated at 37°C for 1 h with 200 μg/ml thermolysin (Sigma, T7902). The reaction was stopped by addition of 400 μM Phosphoramidon (Sigma, R7385) on ice for 20 min. The remaining steps followed AF-03 neutralization of MARV assays.

For AF03-NL neutralization assays, 50 μl/well of serial diluted AF03-NL and AF-03 (starting at 50 μg/ml) were incubated with cells at 37°C for 2 h to enable internalization of the antibodies. 50 μl/well of diluted pseudovirus was then added to a 96-well plate and incubated at 37°C for 36 h. Bright-Glo luciferase reagent was added to detect the relative luciferase intensity. Bioluminescent imaging in vivo

Four-week-old female BALB/c mice were purchased from Beijing Vital River Laboratory Animal Technology Co. Ltd. Mice were intraperitoneally injected with MARV pseudovirus (0.2 ml/mouse). AF-03 (10, 3 or 1 mg/kg) or control antibody Herceptin (10 mg/kg) were injected via intravenous route 24 h and 4 h before the pseudovirus injection respectively. Bioluminescent signals were monitored at Day 4. Briefly, D-luciferin (150 mg/kg body weight) (PerkinElmer, 122799) was intraperitoneally injected into the mice, and then exposed to Isoflurane alkyl for anesthesia. Bioluminescence was measured by the IVIS Lumina Series III Imaging System (Xenogen, Baltimore, MD, USA) with the living Image software. The signals emitted from different regions of the body were measured and presented as average fluxes. All data are presented as mean values ± SEM.

### Evaluation of internalization

HEK293T cells (3×10^5^ cells/well) were washed twice with cold PBS and the supernatant was discarded. AF-03/AF03-NL/human IgG1 isotype (20 μg/ml) was added and incubated at 4°C for 30 min. One group was transferred to 37°C for internalization for 30 min, while the other group continued to be incubated at 4°C for 30 min to adhere to the cell surface. The cells were washed with cold PBS and PE-labeled anti-human IgG Fc secondary antibody (Biolegend, 41070) was added and incubated for 30 min at 4°C. The cells were collected for analysis.

### pHrodo Red labelling and intracellular localization

AF03-NL and AF-03 were covalently labeled with pH-sensitive pHrodo red succinimidyl ester (Thermo, P36600) according to the manufacturer’s instructions. Antibodies were incubated with 10-fold molar excess of pHrodo red succinimidyl ester for 1 h at room temperature. Excess unconjugated dye was removed using PD-10 desalting columns (GE Healthcare). pHrodo Red-labeled antibodies were exchanged into HEPES buffer and concentrated in an Amicon Ultra centrifugal filter unit with a nominal molecular weight cutoff of 30 kDa. Antibody concentration and degree of labeling was determined according to the manufacturer’s instructions.

HEK293T cells (1∼2×10^5^ cells per dish) were cultured overnight in confocal dish pre-treated with polylysine (Beyotime, ST508) and then incubated with pHrodo Red-labeled AF-03 and AF03-NL (20 μg/ml) at 37°C for 30 min. Unbound antibodies were removed by washing with cold PBS. As well, cells were stained with cell membrane dye (DiD) (Thermo, V22887) and Hoechst33342 (Thermo, H1398) at 37°C for 15 min. Single cells were analyzed for pHrodo Red fluorescence on confocal microscopy (dragonfly 200).

### CI-MPR knockin and knockdown

Huh7 Cells were seeded in 6-well plates at 3×10^5^ cells/well and transfected with 2 μg CI-MPR expression plasmids (JetPRIME) when cells reached 40-60% confluence and cultured for 24 h. For CI-MPR silencing, HEK293T cells were seeded in 6-well plates and transfected with siRNA-CI-MPR (Ribbio) and cultured for 48-72 h. CI-MPR expression was detected by flow cytometry.

### Flow cytometry

The cells were harvested and stained with FITC-conjugated anti-CI-MPR antibody (Biolegend, 364207) on ice for 30 min. Cells were washed twice and detected on FACSAria II (BD Biosciences). Data analysis was performed using the FlowJo software.

### Statistical analyses

Data were analyzed, and the graphs were plotted using Prism software (GraphPad Prism 8, San Diego, USA). The data are presented as the mean ± standard error. Statistical differences were compared using unpaired t-tests or ANOVA. The *P* <0.05 was considered statistically significant.

## Results

### Characterization of AF-03

AF-03 was selected from a well-established phage surface display antibody library with immense diversity in the selected complementarity determining region (CDR) loops^32,33^. We further subcloned VH and VL sequences of the antibody into a mammalian full-length immunoglobulin expression vector for full-length IgG expression. As shown in Fig. 1A, AF-03 was eukaryotically expressed with the purity over 95%. To determine the binding affinity, recombinant MARV GP without MLD was prepared (Fig. 1A). ELISA analysis showed the remarkable binding of AF-03 to MARV GP (Fig. 1B). Furthermore, SPR assay was done to determine the binding kinetics and showed that AF-03 bound to MARV GP with high affinity (K_D_ value was 4.71×10^-11^M) (Fig. 1C). To identify determinants of MARV GP binding to AF-03, we utilized computer-guided homology modeling and molecular docking to generate computer models of MARV GP in complex with AF-03. Specifically, we obtained the theoretical 3-D structure of AF-03 Fv (Fig. 1D). Based on the 3-D structure of AF-03 and MARV GP separately, the 3-D complex structure of AF-03 and MARV GP achieved utilizing the molecular docking method (Fig. 1E). Overall, these data suggest the potency of AF-03 binding to MARV GP.

**Figure 1.**
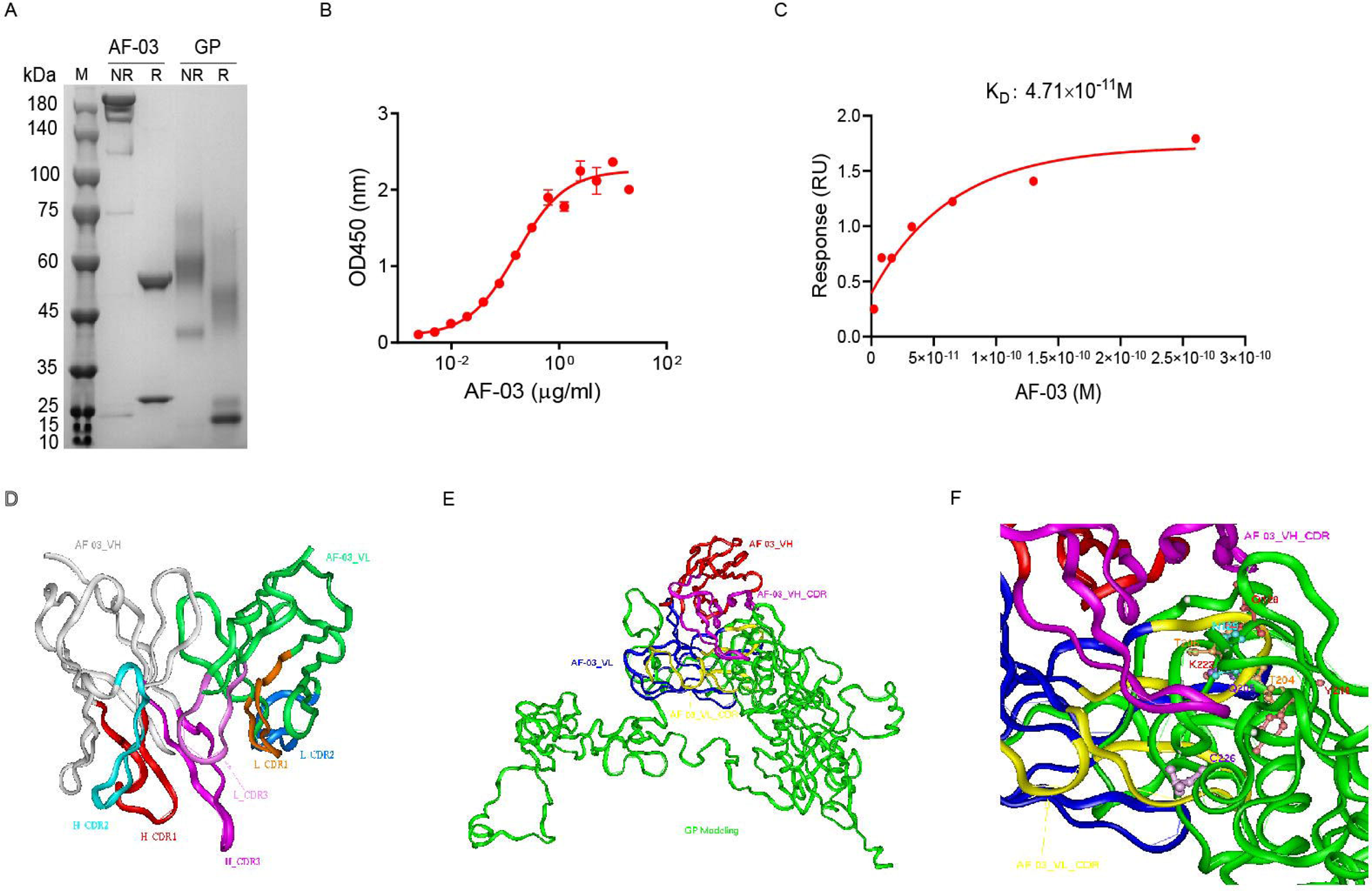
Binding activity of mAb AF-03 to MARV GP and its epitopes. (A) AF-03 and MARV GP proteins are examined by SDS-PAGE. NR, non-reducing; R, reducing. (B) The binding capacity of AF-03 to MARV GP is determined by ELISA. The absorbance is detected at 450 nm. (C) The binding kinetics of AF-03 to MARV GP is detected by SPR assay. Experiments are independently repeated at least three times, and the data from one representative experiment is shown. (D) The 3-D ribbon structures of the AF-03 Fv fragment. The red ribbon denotes H-CDR1, the light blue denotes H-CDR2, the pink denotes H-CDR3, the orange denotes L-CDR1, the deep blue denotes L-CDR2, the purple denotes L-CDR3. (E) AF-03 and MARV GP complex derived from theoretical modeling. The green ribbon denotes the orientation of the MARV GP fragment, the yellow denotes AF-03 VLCDR, the pink denotes AF-03 VHCDR, the deep blue denotes AF-03 VL and the red ribbon denotes AF-03 VH. (F) By molecular docking analysis of van der Waals interaction, intermolecular hydrogen bonding, polarity interaction and electrostatic interaction, the key amino acid residues of MARV GP are screened.

### Epitope mapping of MARV GP bound to AF-03

Under CVFF forcefield, chosen steepest descent and conjugate gradient minimization methods, after 30,000 steps of minimization with the convergence criterion 0.02 kCal/mol, the optimized structure of the AF-03 Fv was evaluated (Fig.1D). Using a Ramachandran plot, the assignment of the whole heavy atoms of the AF-03 Fv was in the credible range. Using the crystal complex structure of MR78 and GP protein as model, based on the optimized theoretical 3-D structure of MARV GP protein, the theoretical 3-D complex structure of AF-03 and MARV GP protein was constructed (Fig.1E). Through analyzing the van der Waals interactions, inter-molecular hydrogen bonds, polar interactions and electrostatic interactions between AF-03 and MARV GP, defining the binding region distance towards CDRs of AF-03 Fv fragment as 0.6 nm, the key amino acid residues of MARV GP were predicted for amino acid point mutations (Fig. 1F). Based on the volume and character of amino acid residues, the residues T^204^–Q^205^–T^206^, Y^218^ and K^222^ were mutated to alanine, and Q^128^–N^129^ were mutated to serine as well as C^226^ was mutated to tyrosine. Firstly, we investigated if this mutated MARV species was still sensitive to AF-03 treatment. The inhibition assay revealed the significant impairment of neutralizing activity of AF-03 to MARV pseudovirus harboring Q^128^S-N^129^S or C^226^Y compared with WT MARV and those loading other indicated mutations (Fig. 2A left panel), which indicates that Q^128^/N^129^/C^226^ functions as key amino acids responsible for AF-03 neutralization. Furthermore, we constructed the mutated MARV GP. ELISA assay showed that Q^128^S-N^129^S or C^226^Y mutation significantly disrupted the binding of GP to AF-03, while the binding and neutralizing capacity of MR78 to mutant GP and pseudovirus harboring C^226^Y instead of Q^128^S-N^129^S, a mAb reported to be isolated from Marburg virus survivors^16^, was comparable to WT counterparts (Fig. 2A right panel and B). Furthermore, we analyzed the secondary structure of the MARV GP and its mutants. By circular dichroism, the structure of both mutants was not obviously different from that of the parental GP (Fig. 2C, Tab. s1). Therefore, the weakened binding to the antibody was not due to the conformational change of GP caused by the mutation. Competitive ELISA showed that AF-03 and MR78 could compete with each other to bind to MARV GP (Fig. 2D). Collectively, these results indicate the epitopes of these two mAbs overlapped partially.

**Figure 2.**
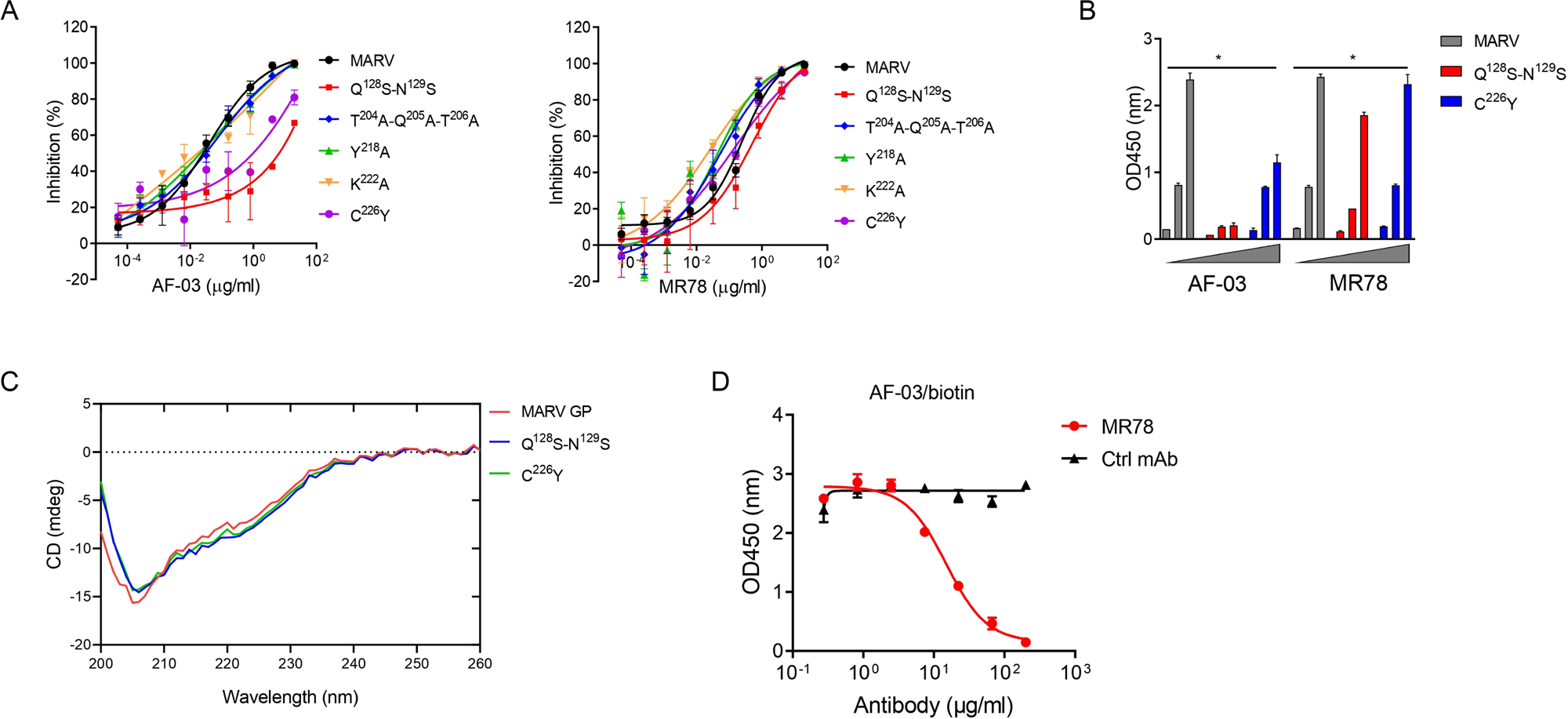
AF-03 Epitope Identification. (A) The neutralization activity of AF-03 or MR78 to mutated pseudovirus (Q^128^S-N^129^S, Q^204^A-T^205^A-Q^206^A, Y^218^A, K^222^A, C^226^Y) is evaluated in HEK293T cells. The inhibition rate is analyzed. (B) The binding of AF-03 and MR78 to mutant GP (Q^128^S-N^129^S or C^226^Y) is examined by ELISA respectively. (C) Secondary structure of MARV GP and mutants are detected by CD. (D) The epitope overlapping between AF-03 and MR78 is examined by the competitive ELISA.

### In vitro neutralizing activity of AF-03

Given the high binding affinity of AF-03 to MARV GP, we sought to determine whether AF-03 could impede pseudotyped MARV viral entry. An in vitro neutralization assay was developed based on full-length MARV GP-pseudotyped virus using a HIV vector (pSG3.Δenv.cmvFluc). Liver and adrenal glands have been reported to be the early targets of MARV infection ^14,35^. Therefore, we first tested the entry of MARV to hepatocyte cell line (Huh7) and renal cell line (HEK293T cells) by measuring the relative luciferase intensity. These two cell lines were susceptible to MARV cell entry. We used MR78 and cetuximab as positive and negative controls respectively. As expected, cetuximab had no effects on pseutotyped MARV entry. In contrast, AF-03 actively inhibited viral entry to HEK293T cells, with IC_50_ value of 0.13 μg/ml. As well, IC_50_ of MR78 was 0.44 μg/ml (Fig. 3A left panel). In Huh7 cells, IC_50_ of AF-03 and MR78 was 0.4 and 1.03 μg/ml, respectively (Fig. 3A right panel). These results suggest that AF-03 has stronger potency of neutralization than MR78. We also conducted AF-03 neutralization experiments on pseudotyped Angola, Musoke and RAVN strains and showed strong and comparable neutralizing ability to all these strains (IC_50_ was 0.32, 0.12 and 0.15 μg/ml respectively) (Fig. 3B). Taken together, these data suggest that AF-03 harbors prominent neutralizing activity to MARV infection.

**Figure 3.**
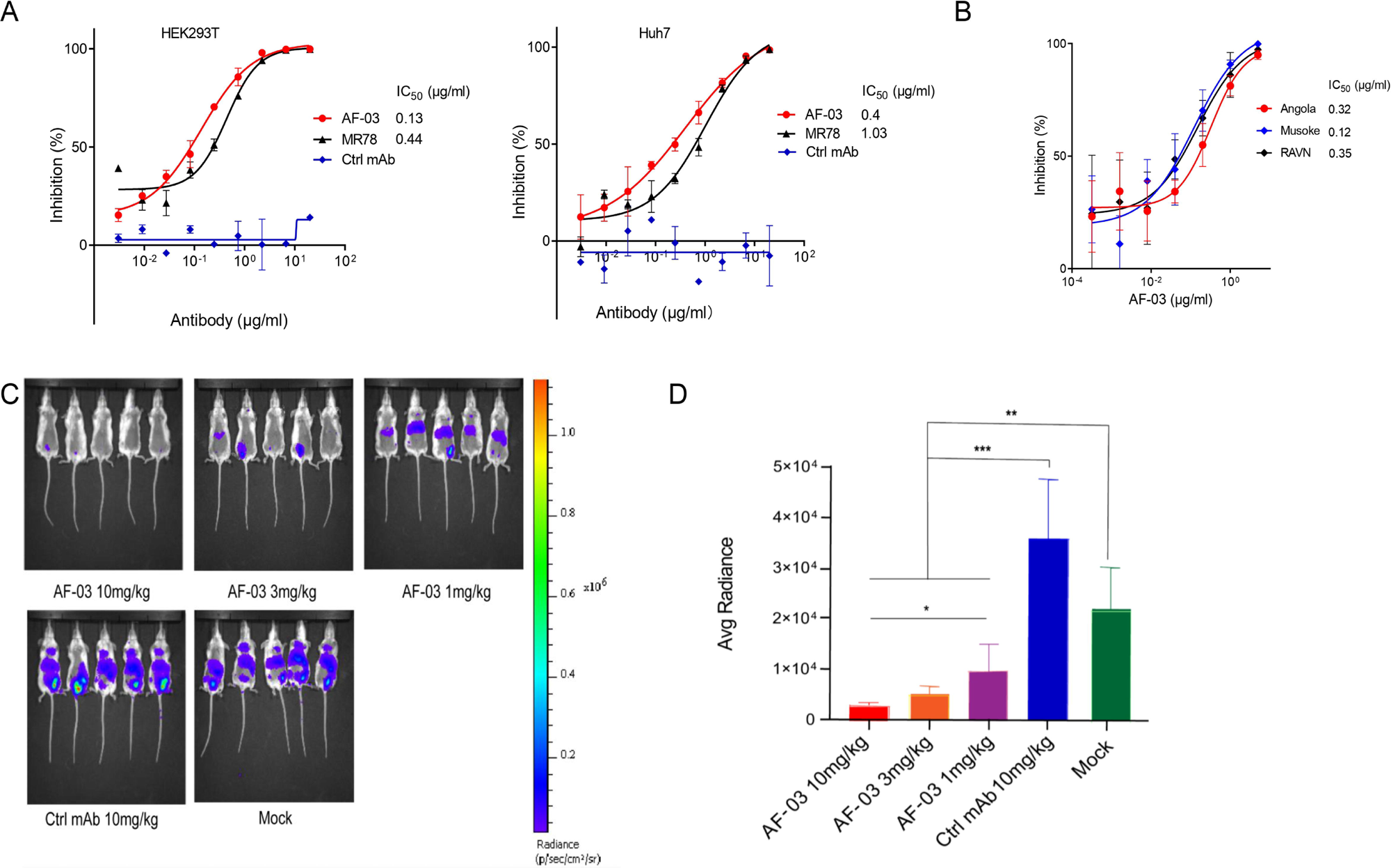
In vitro and in vivo neutralization of MARV pseudovirus infection by AF-03. (A) Pseudotypic MARV-Uganda is incubated with AF-03, MR78 or control mAb at 37°C for 1 h before infecting HEK293T cells (left) and Huh7 cells (right) respectively. Luciferase is assayed and inhibition rates are calculated. (B) Pseudotypic MARV-Angola, Musoke and RAVN infect HEK293T cells respectively and neutralization activity of AF-03 to these species is determined. (C) AF-03 (10, 3, 1mg/kg) is administrated at 24 and 4 h before intraperitoneal injection of MARV pseudovirus. On day 4, bioluminescence signals are detected by an IVIS Lumina Series III imaging system. (D) The average radiance value based on the luminescence of (C). *\*p<0.05, **p<0.01, ***p<0.001*.

### In vivo preventive efficacy of AF-03

To verify the in vivo preventive efficacy, AF-03 was intravenously injected into mice before pseudotyped MARV exposure (-24 h and -4 h) respectively. The bioluminescence intensity was measured on day 4 after pseudovirus injection. As shown in Fig. 3C and D, AF-03-treated group displayed lower bioluminescence activity compared with the control group, while the treatment with control antibodies had no effects. Administration of 1 mg/kg AF-03 prior to the injection of MARV could decrease viral infection to approximately 50% level and increasing doses of AF-03 led to higher preventive efficacy. This indicates clearly that AF-03 is capable of preventing from MARV infection in a dose-dependent manner and represents a potential candidate for MARV treatment.

### AF-03 impedes cell entry of EBOV, SUDV and BDBV harboring GPcl

Given the close structural similarity of Marburg virus to ebola virus, to determine whether AF-03 was also available to the treatment of EBOV infection, we conducted neutralization of pseudotyped EBOV, SUDV and BDBV by AF-03. In addition, considering that glycan cap or mucin-like domain are known to mask the putative receptor-binding domain on EBOV, SUDV and BDBV GP and thus impede the engagement between AF-03 and GP^16^, glycan cap and mucin-like domain were enzymatically cleaved by digesting GP with 250 μg/ml thermolysin at 37°C. The infection of pseudotyped ebolavirus harboring cleaved GP to host cells was comparable or stronger than those containing intact GP (Fig. s1). Intriguingly, the inhibitory function of AF-03 on cell entry of all three species of ebolavirus bearing cleaved GP was much stronger than those bearing uncleaved GP (Fig. 4), which suggests that AF-03 has therapeutic potentials for EBOV, SUDV and BDBV infection.

**Figure 4.**
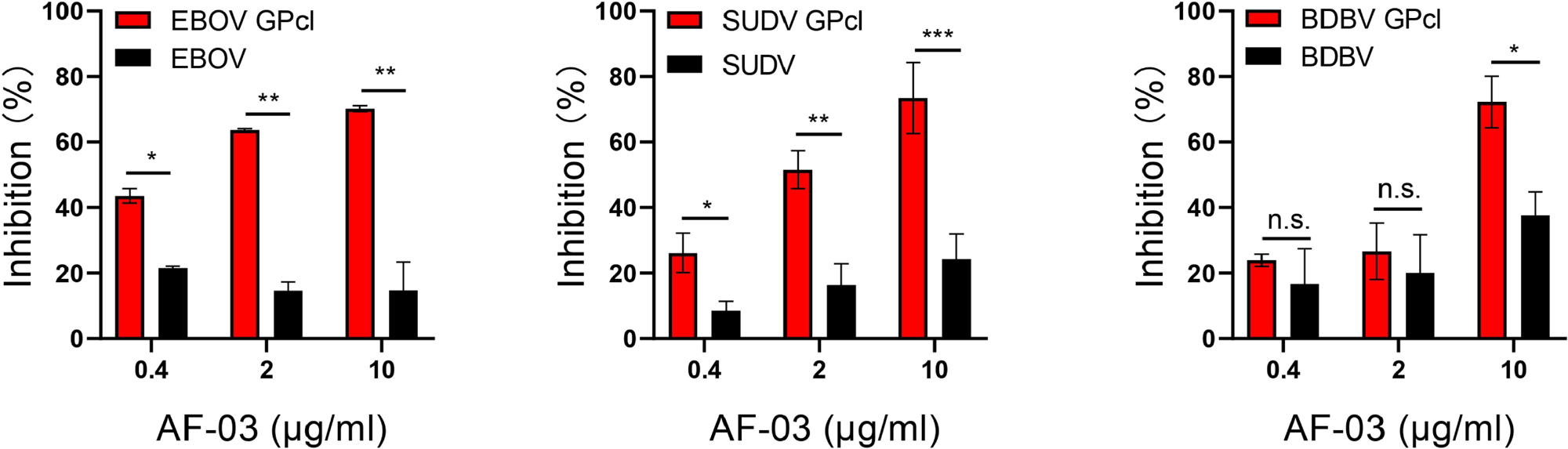
The neutralization activity of AF-03 to EBOV, SUDV and BDBV harboring cleaved GP. Pseudotypic EBOV, SUDV, and BDBV are processed with thermolysin at 37°C. Inhibition of these ebola virus entry harboring GP or GPcl by AF-03 is examined by luciferase assay. *\*p<0.05, **p<0.01, ***p<0.001*.

### The potency of NPC2-fused AF-03 to be delivered into the endosome

Given the inability of AF-03 to transport into endosomal compartment where intact GP is cleaved by cathepsin B/L, we engaged NPC2 to the N-terminus of light chain of AF-03 (Fig. s2A), according to a protocol described previously^36^. As well, we produced the 1-3 domain of CI-MPR (Fig. s2A), which is a ligand for NPC2 and expressed on the cellular and endosomal membrane^37^. The results showed that NPC2-fused AF-03 (termed AF03-NL), rather than AF-03, bound to CI-MPR1-3 (Fig. s2B). Next, we investigated the internalization of AF03-NL. AF03-NL or AF-03 was incubated with HEK293T cells, which expressed CI-MPR (Fig. s3A), at 4L for attachment. As expected, AF03-NL instead of AF-03 adhered to the cell surface, detected by fluorescence-labelled secondary IgG. Upon endocytosis, the fluorescence on the cell surface decreased dramatically, implying the occurrence of AF03-NL internalization (Fig. 5A). To further address this issue, AF-03 and AF03-NL were labelled by pHrodo Red dye that is sensitive to acidic niche. Consequently, pHrodo Red-conjugated AF03-NL was observed in the acidic endosomal compartment by flow cytometry and fluorescence microscopy respectively (Fig. 5B and C). In contrast, AF-03 was not seen in the endosome.

**Figure 5.**
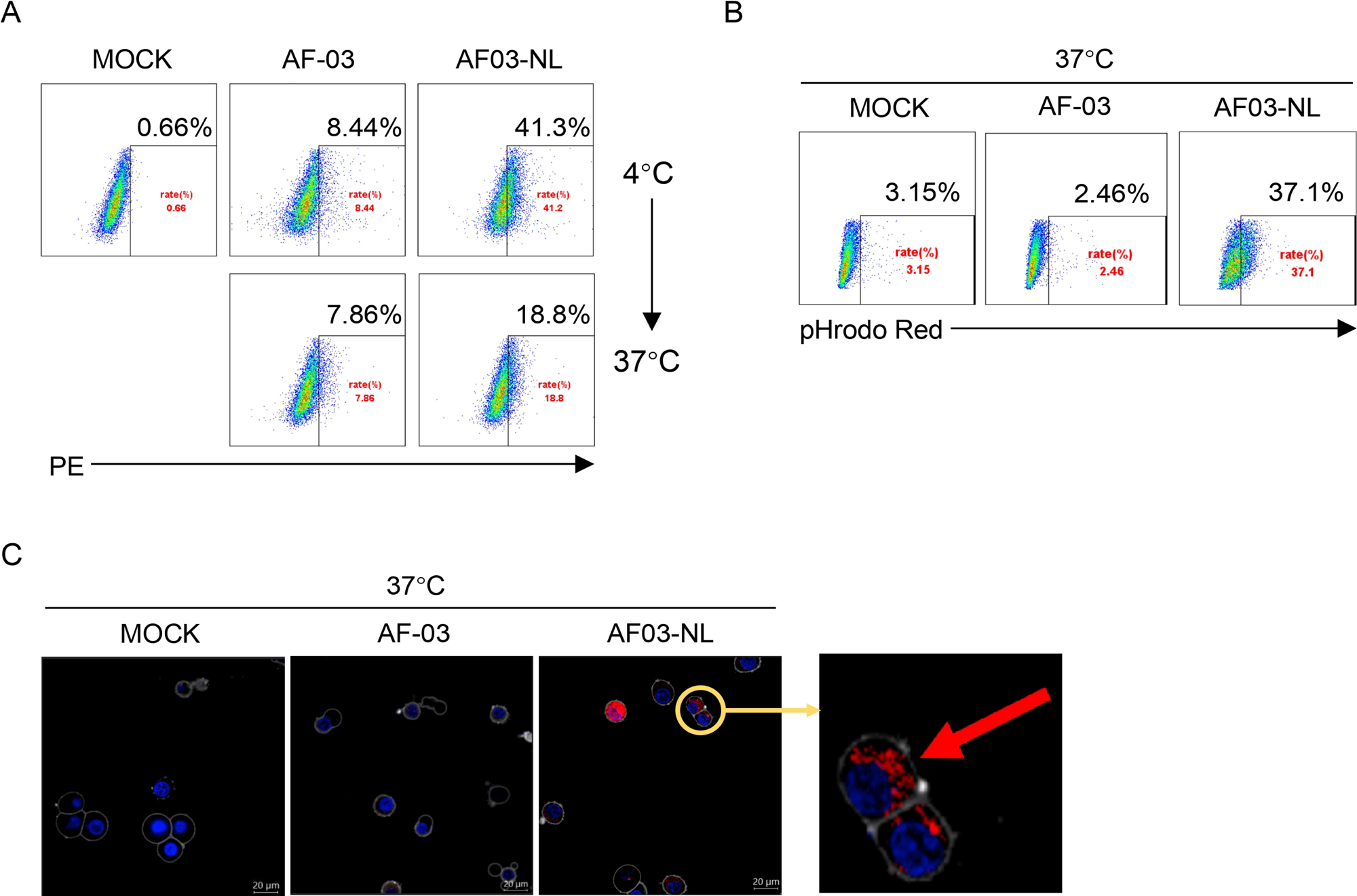
Cellular internalization of AF03-NL. (A) AF-03, AF03-NL or human IgG1 isotype (MOCK) is incubated with cells at 4L for 1 h to prevent internalization and then at 37L for another 2 h to allow internalization. PE-conjugated secondary antibody is added prior to analysis by flow cytometry. (B,C) pHrodo Red-labelled AF-03 or AF03-NL is incubated with cells at 37□ for 1 h and analyzed by flow cytometry (B) and fluorescence microscopy (C) respectively. The red arrow denotes internalized AF03-NL. Experiments are independently repeated at least three times, and the data from one representative experiment is shown.

### Pan-filovirus inhibition of cell entry by AF03-NL via engagement between NPC2 and CI-MPR

Firstly, we compared the binding of AF03-NL/AF-03 to MARV GP. ELISA showed relatively weak binding activity of AF03-NL compared with AF-03 (Fig. s4A). We thereafter evaluated the neutralizing activity of AF03-NL/AF-03 to MARV pseudovirus. Intriguingly, AF03-NL showed stronger neutralizing activity than AF-03 (The IC_50_ was 0.057 and 0.284 μg/ml, respectively) (Fig. s4B), which may be attributed to sustained tethering of AF03-NL to pseudovirus at extracellular space as well as endosomal compartment. Secondly, we compared the neutralizing activity of AF03-NL and AF-03 to a series of filovirus species. AF03-NL displayed superior neutralizing activity to other nine filovirus species. While, no or weak inhibition of entry by AF-03 was found (Fig. 6A). Furthermore, AF03-NL, instead of AF-03, also actively inhibited cell entry of 17 EBOV mutants that were detected previously in natural hosts (Fig. 6B).

**Figure 6.**
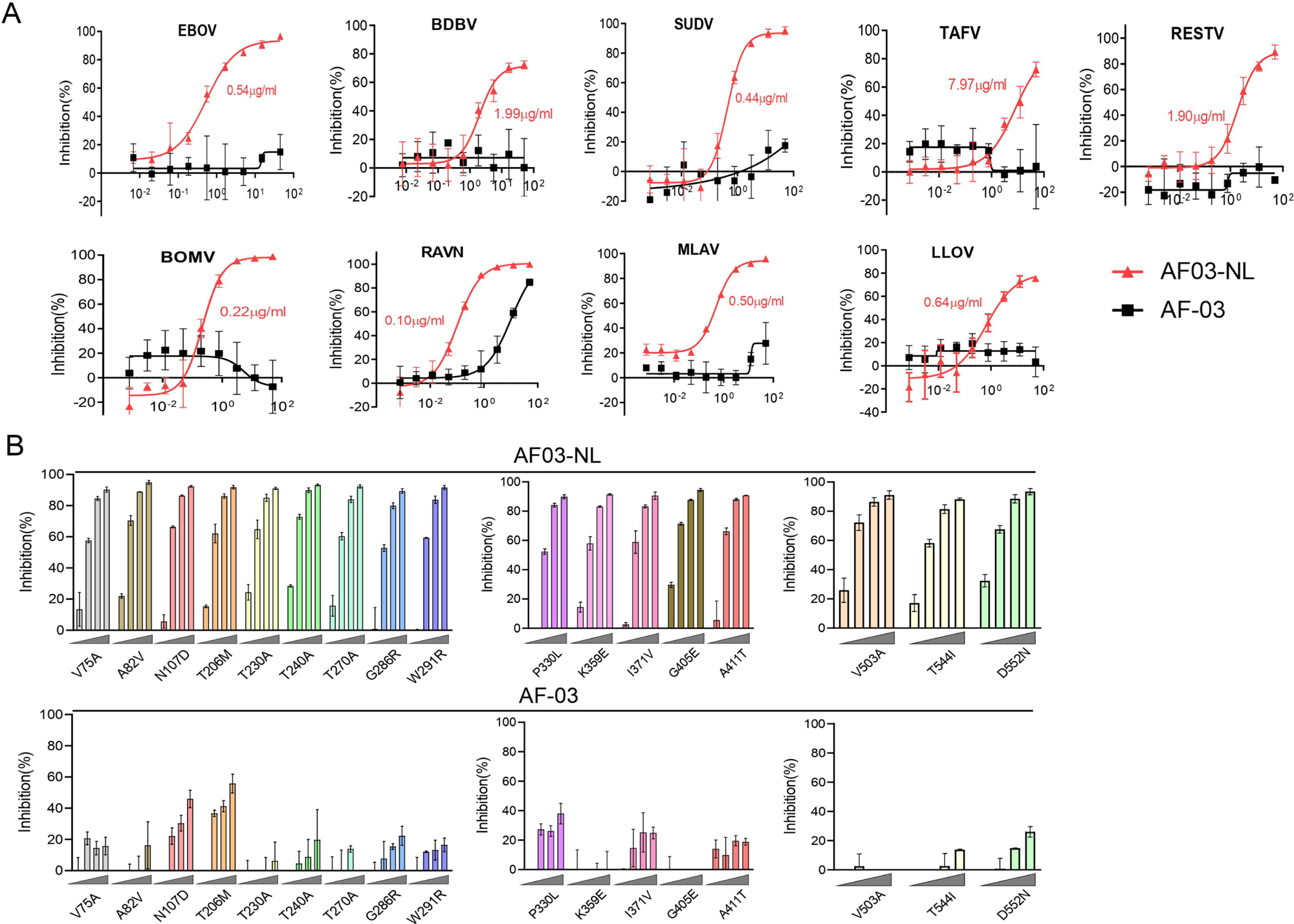
Pan-filovirus entry inhibition by AF03-NL. (A,B) AF-03 or AF03-NL (50-0.0007 μg/ml, 4-fold dilution) is incubated with HEK293T cells at 37°C for 2 h prior to exposure to pseudotypic filovirus species (A) and EBOV mutants (B). Luciferase is assayed and inhibition rates are calculated.

To investigate the mechanisms underlying the potency of AF03-NL, we produced NPC2 protein (Fig.7A) and then examined the inhibition of EBOV entry by AF03-NL or the mixture of AF-03 and NPC2. AF-03, NPC2 alone or in combination did not inhibit EBOV entry. Conversely, AF03-NL actively impeded this process (Fig. 7B). To clarify whether this effect is CI-MPR-dependent, CI-MPR in HEK293T cells was silenced (Fig. 7C). The result showed that CI-MPR knockdown rendered significant abrogation of the inhibitory ability of AF03-NL (Fig. 7C). We also introduced CI-MPR into Huh7 cell line that is null for this receptor (Fig. s3B). The inhibitory effects of AF03-NL were augmented in CI-MPR-overexpressed cells compared with empty vector-introduced counterparts (Fig. 7D). Taken together, these data indicate that the inhibitory potency of AF03-NL is dependent on the interaction between NPC2 and CI-MPR.

**Figure 7.**
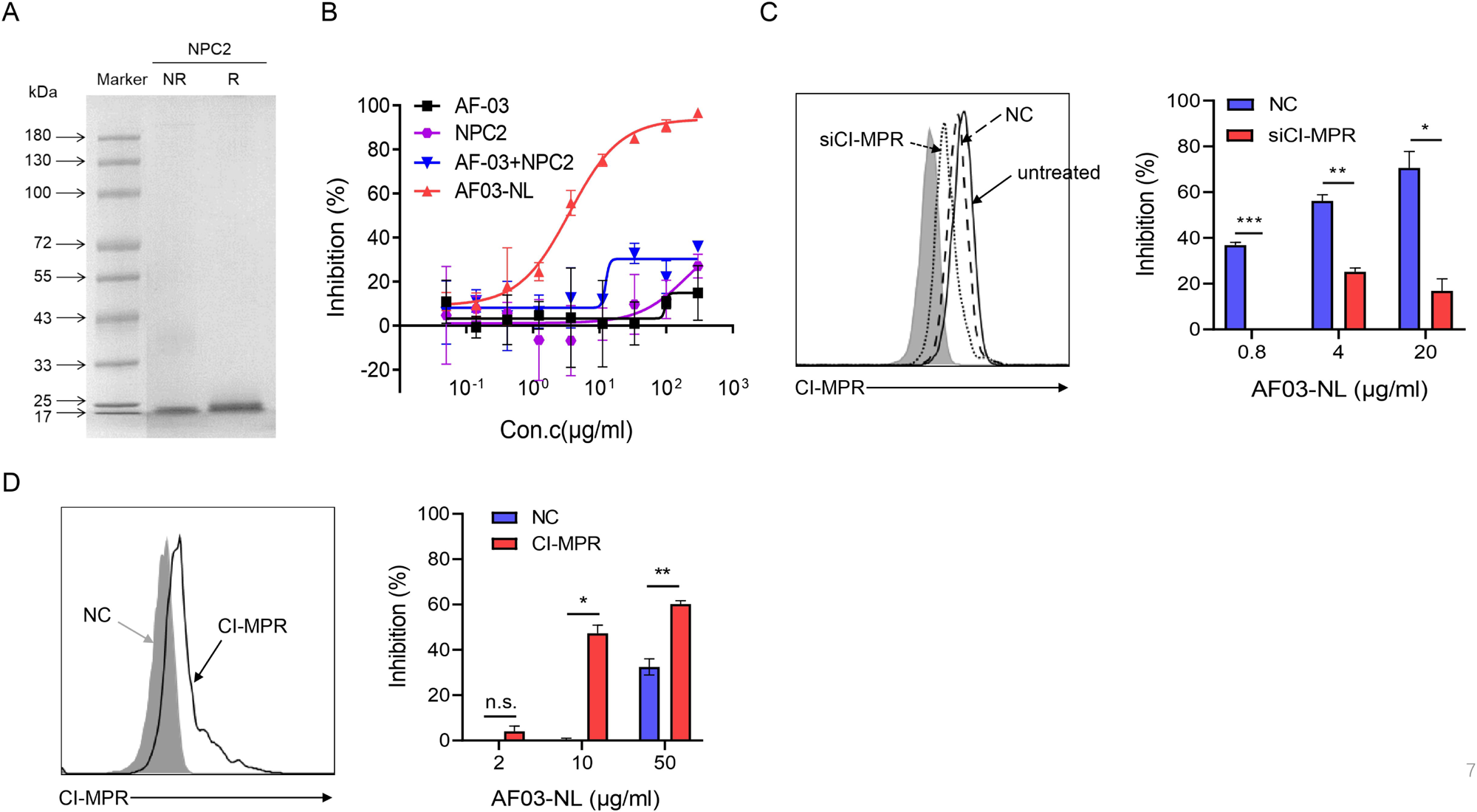
The requirement of CI-MPR for the neutralization activity of AF03-NL. (A) NPC2 protein is examined by SDS-PAGE. NR, non-reducing; R, reducing. (B) AF03-NL, AF-03, NPC2 alone or equimolar combination of AF-03 and NPC2 is incubated with HEK293T cells at 37°C for 2 h prior to exposure to pseudotypic EBOV. Luciferase is assayed and inhibition rates are calculated. (C) HEK293T cells are treated with siRNA-CI-MPR or negative control vector (NC) respectively and CI-MPR expression is detected by flow cytometry. AF03-NL is incubated with siCI-MPR or NC-treated HEK293T cells at 37°C for 2 h respectively prior to exposure to pseudotypic EBOV. (D) CI-MPR is introduced into Huh7 cells and its expression is detected by flow cytometry. AF03-NL is incubated with CI-MPR or NC-knockin Huh7 cells at 37°C for 2 h respectively prior to exposure to pseudotypic EBOV. Luciferase is assayed and inhibition rates are calculated.

## Discussion

The Marburg virus was initially identified after simultaneous outbreaks in Marburg and Frankfurt in Germany in 1967^16,38^. To date, there have been a dozen outbreaks of Marburg virus infection in humans^39^. Giving the recurrence of Marburg virus outbreaks and its high virulence and lethality, there is an urgent need to develop prophylactic and therapeutic interventions for Marburg infections. MARV GP is a surface viral protein, which is responsible for host receptor binding and cell entry thus provides an attractive target for the development of antagonists. Flyak et al. screened several MARV GP-specific neutralizing antibodies from the PBMC samples of a MARV-infected survivor, which achieved 100% protection in mice subjected to mouse-adapted MARV challenge^29^. The MARV GP-specific antibody cocktail was also developed, three mAbs cocktail could protect hamster from lethal hamster-adapted MARV infection, while treatment with either one or two antibodies failed^40^.

In this study, we selected an antibody from a human antibody phage library and the affinity constant reached the 10^-11^M level. The neutralizing activities of the antibody were demonstrated by utilizing pseudotyped MARV Uganda strain. The results showed that AF-03 effectively inhibited HIV vector (pSG3.Δenv.cmvFluc) pseudotyped MARV viral entry at IC_50_ of 0.13 and 0.4 μg/ml in HEK293T and Huh7 respectively. Furthermore, compared with control antibody, AF-03 exhibited a protective property against pseudovirus infection in mice. Epitope mapping results showed that Q^128^–N^129^ and C^226^ of GP was the binding and functional epitopes that interacted with AF-03, which means AF-03 targeting the interface of GP-NPC1 interaction, considering that N^129^ is known to be located in the NPC1 binding domain. RBD is highly conserved among filovirus species, so it is an attractive target for broadly effective anti-filovirus drug development^41^. We found that AF-03 also bound to EBOV GPcl and could neutralize ebola viruses bearing cleaved GP in vitro, suggesting that AF-03 represents a good candidate for endosome-delivering strategy by ligation to another mAb against a surface-exposed EBOV GP epitope or a ligand peptide for host cation-independent mannose-6-phosphate receptor^36^, which will ultimately afford cross-reactivity against multiple filovirus species. Accordingly, we designed NPC2-fused AF-03 and demonstrated its broad-spectrum inhibitory capacity to filovirus species and EBOV mutants. Future investigations on the inhibition of AF03-NL to authentic virus infection in vitro and in vivo are warranted.

Overall, our study identified a high-affinity anti-MARV antibody AF-03 targeting a conserved and hidden site at the filovirus GPcl-NPC1 interface, which was capable of neutralizing MARV infection both in vitro and in vivo. Furthermore, AF-03 may be a potential candidate for the effective protection against pan-filovirus species infection. Investigations on AF-03 treatment of mice challenged by authentic virus are undergoing.

## Abbreviations

MARV: Marburg virus
EBOV: Ebola virus
SUDV: Sudan virus
BDBV: Bundibugy virus
mAb: monoclonal antibody
PBS: phosphate-buffered saline
GP: glycoprotein
RAVN: Ravn virus
RESTV: Reston virus
TAFV: Tai forest virus
MVD: Marburg virus disease
EVD: EBOV virus
RBD: receptor binding domain
MLD: mucin like domain
NPC1: Niemann-Pick C1
CDR: complementarity determining region
CVFF: consistent valence force field
FV: fragment variable
CI-MPR: cation-independent mannose-6-phosphate receptor
NPC2: Niemann-Pick C2
CD: circular dichroism

## Disclosure of interests

Yuting Zhang, Min Zhang, Haiyan Wu, Xinwei Wang, Hang Zheng, Junjuan Feng, Jing Wang, Longlong Luo, He Xiao, Chunxia Qiao, Xinying Li, Yuanqiang Zheng, Weijin Huang, Youchun Wang, Yi Wang, Yanchun Shi, Jiannan Feng, Guojiang Chen declare that they have no competing interests.

## Author contributions

Guojiang Chen, Jiannan Feng, Yanchun Shi and Yi Wang conceived and designed this study. Guojiang Chen, Jiannan Feng and Yuanqiang Zheng provided funding support. Yuting Zhang, Min Zhang, Haiyan Wu and Xinwei Wang performed of the experiments and prepared the manuscript. Hang Zheng and Junjuan Feng were involved in optimization of the experimental protocols. Jing Wang, Longlong Luo, He Xiao, Chunxia Qiao, Xinying Li, Yuanqiang Zheng, Weijin Huang, Youchun Wang provided methodological support. All authors contributed to the article and approved the submitted version.

## Acknowledgments

This work is granted from the National Natural Science Foundation of China (81672803, 31771010, 81871252).

## Ethics approval

The animal study was reviewed and approved by the Institutional Animal Care and Use Committee of Academy of Military Medical Sciences.

## Notes

### Competing Interest Statement

The authors have declared no competing interest.

### Summary of Updates

Figure 1C-F,2A-B,3C,4 revised; Supplemental files updated; Section on materials and methods updated to clarify SPR analysis of antibody affinity, computer‑guided homology modeling and molecular docking.

