## Supplemental files for "A novel MARV glycoprotein-specific antibody with potentials of broad-spectrum neutralization to filovirus"

**Supplementary Table**

Tab.s1 Proportion of secondary structure of parent and mutant MARV GP

|  | MARV GP | Q^128^S-N^129^S | C^226^Y |
| --- | --- | --- | --- |
| Helix | 18.15% | 19.46% | 19.28% |
| Antiparallel | 13.66% | 12.95% | 13.03% |
| Parallel | 12.10% | 11.78% | 11.85% |
| Beta-Turn | 18.61% | 18.44% | 18.39% |
| Rndm. Coil | 37.49% | 37.37% | 37.45% |
| Total Sum | 100.00% | 100.00% | 100.00% |

The mutant protein and the parent MARV GP were purified and replaced by PBS ultra-filtration. The secondary structure of the protein was analyzed by Circular Dichroism.

**Supplementary Figure legends**

**Fig. s1. Comparison of the cellular entry capacity of ebolavirus harboring cleaved or intact GP.**

Ebola virus (EBOV, SUDV, BDBV) were treated with or without thermolysin at 37°C and then infected HEK293T cells. Luciferase intensity was assayed. **p<0.05, **p<0.01.*

**Fig. s2. Characterization of AF03-NL and CI-MPR1-3.**

(A) AF03-NL and CI-MPR1-3 domain proteins are examined by SDS-PAGE. NR, non-reducing; R, reducing. The arrow denotes NPC2-fused light chain. (B) The binding capacity of AF03-NL and AF-03 to CI-MPR1-3 is detected by ELISA.

**Fig. s3. CI-MPR expression in HEK293T and Huh7 cells.**

The CI-MPR expression in (A) HEK293T and (B) Huh7 cells is examined by flow cytometry.

**Fig. s4. Comparable binding and inhibitory activity of AF03-NL and AF-03.**

(A) The binding capacity of AF-03 and AF03-NL to MARV GP is detected by ELISA. (B) AF-03 orAF03-NL is incubated with HEK293T cells at 37°C for 2h prior to exposure to pseudotypic MARV-Uganda. Luciferase is assayed and inhibition rates are calculated.

**Fig. s1**

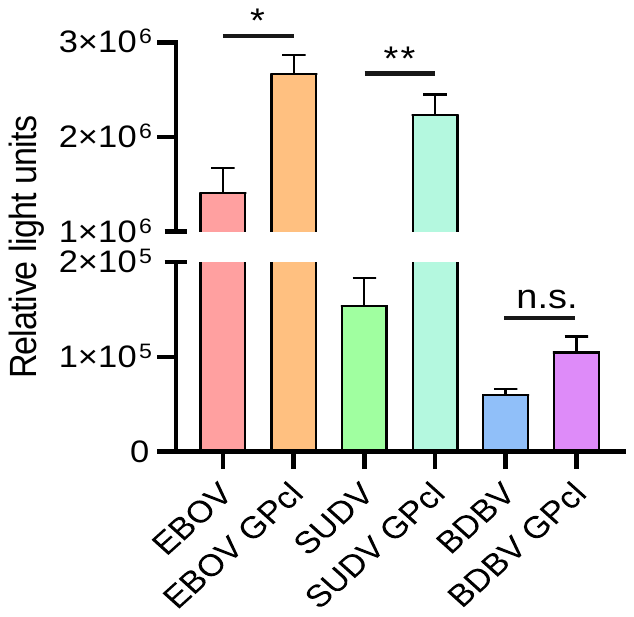

**Fig. s2**

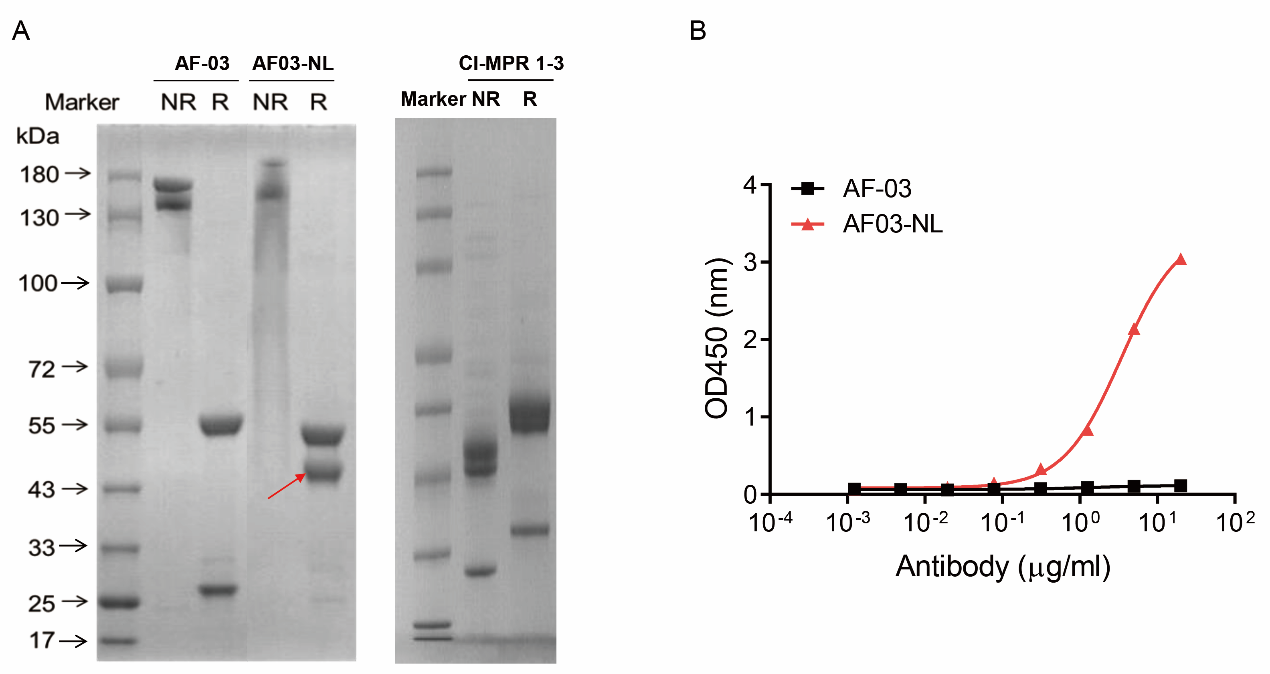

**Fig. s3**

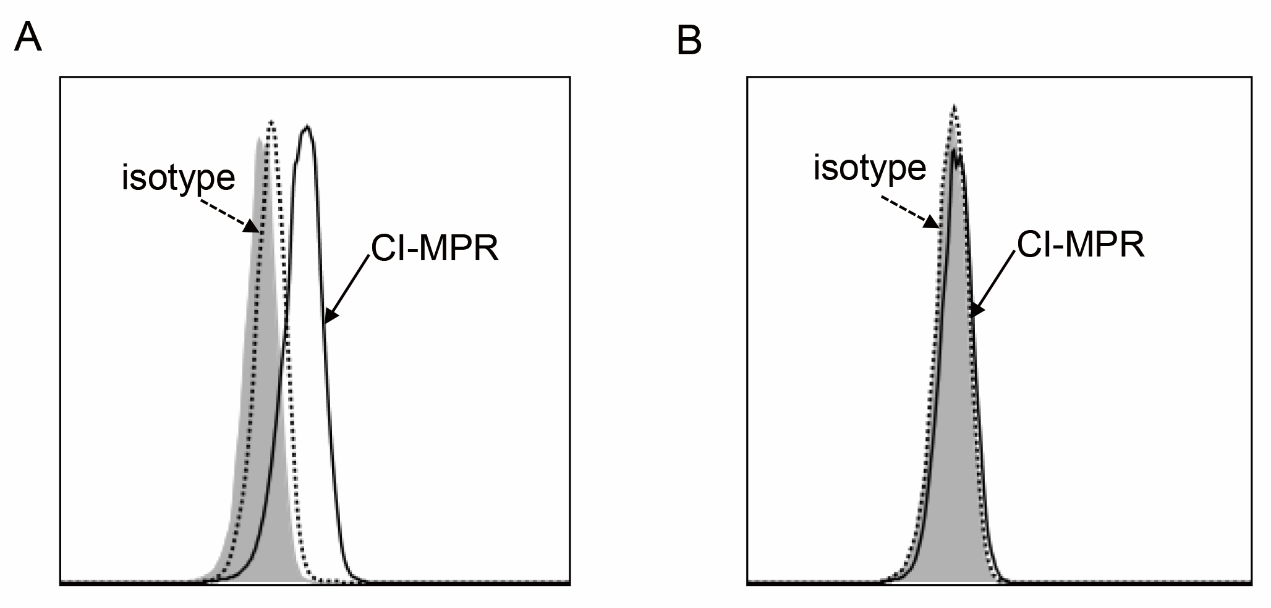

**Fig. s4**

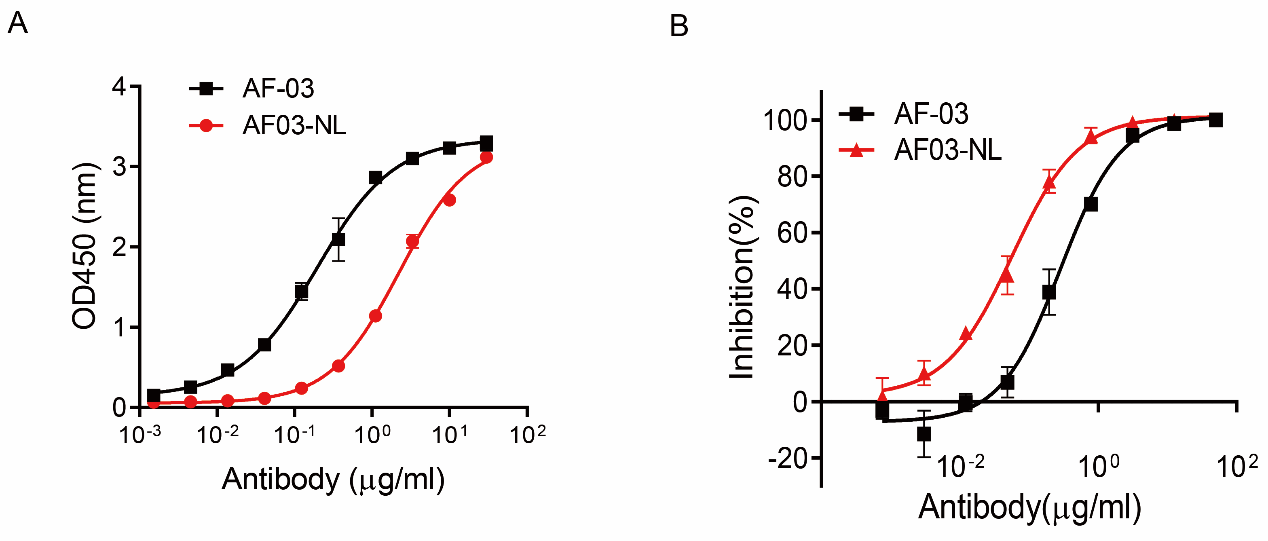
